## Supplemental Figures for "Computational and cellular studies reveal structural destabilization and degradation of MLH1 variants in Lynch syndrome"

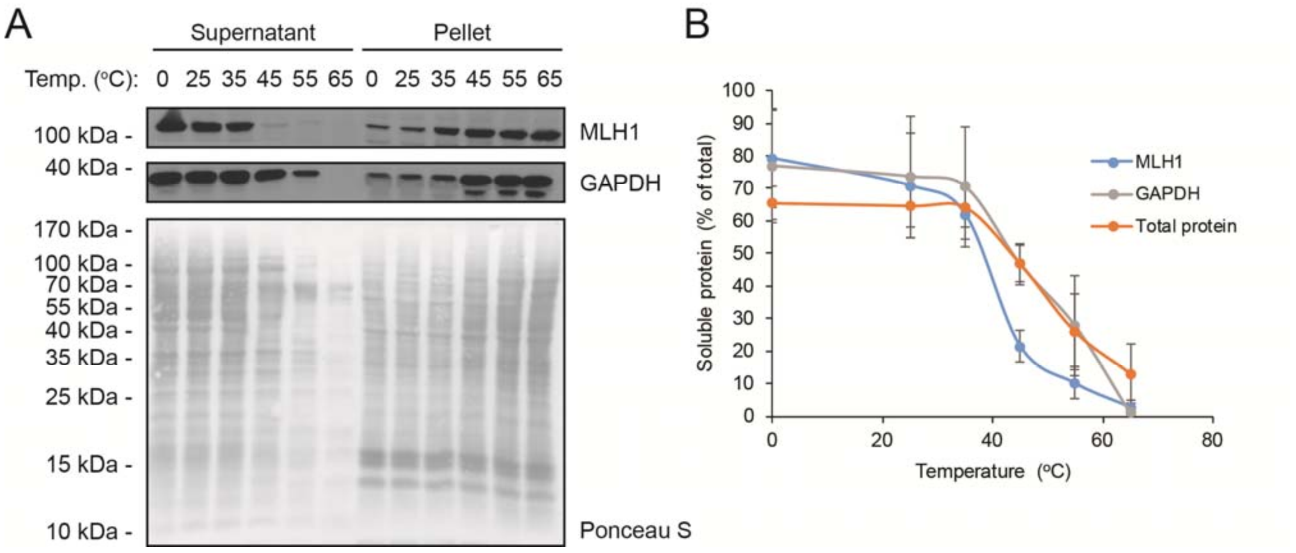


**Figure supplement – Figure 2.1** – *Solubility of wild-type MLH1 at a range of temperatures.* (A) Transfected cells were lysed by sonication and incubated for 30 min. at the indicated temperatures. Then the lysates were separated into supernatant and pellet fractions by centrifugation. Western blotting using antibodies to MLH1 was employed to determine the amount of MLH1 in the fractions. Ponceau S staining and blotting with antibodies to GAPDH served as controls. (B) Quantification of blots as shown in panel (A) showing the amount of soluble protein (supernatant) normalized to total (supernatant + pellet). The total protein quantification is based on the full lanes of the Ponceau S stainings. The error bars indicate the standard deviation (n=3).

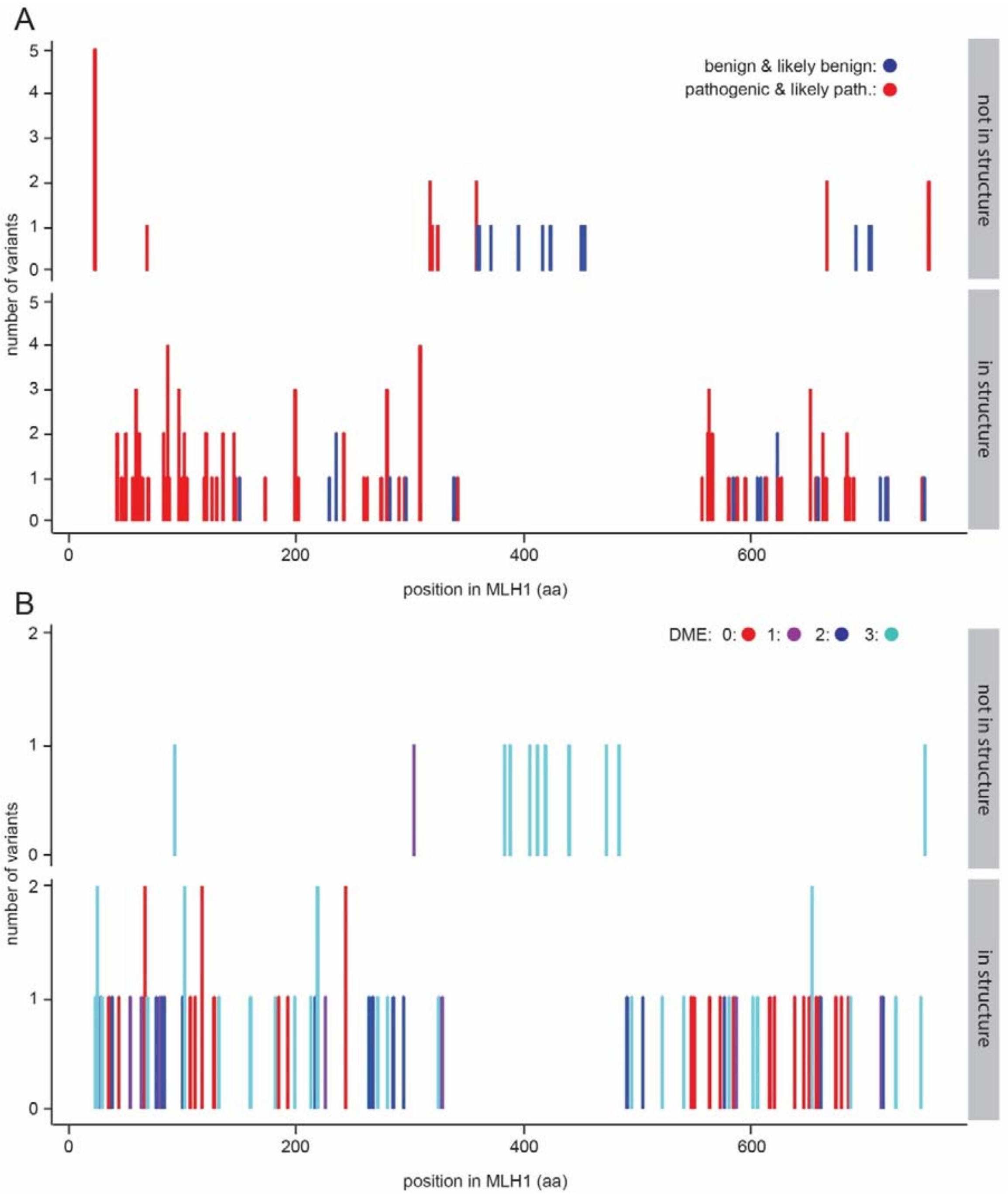


**Figure supplement – Figure 2.2** – *Variants within the central disordered region.* (A) ClinVar benign and likely benign variants (blue) are found throughout the protein, while pathogenic and likely pathogenic variants (red) are enriched in the structured domains. (B) The variants previously tested for function (Takahashi et al., 2007) are distributed throughout the MLH1 protein. Similar to the observations from ClinVar, functional variants (DME score 3) are seen throughout the protein, while loss-of-function in multiple assays (DME scores <2) are enriched in the structured domains.

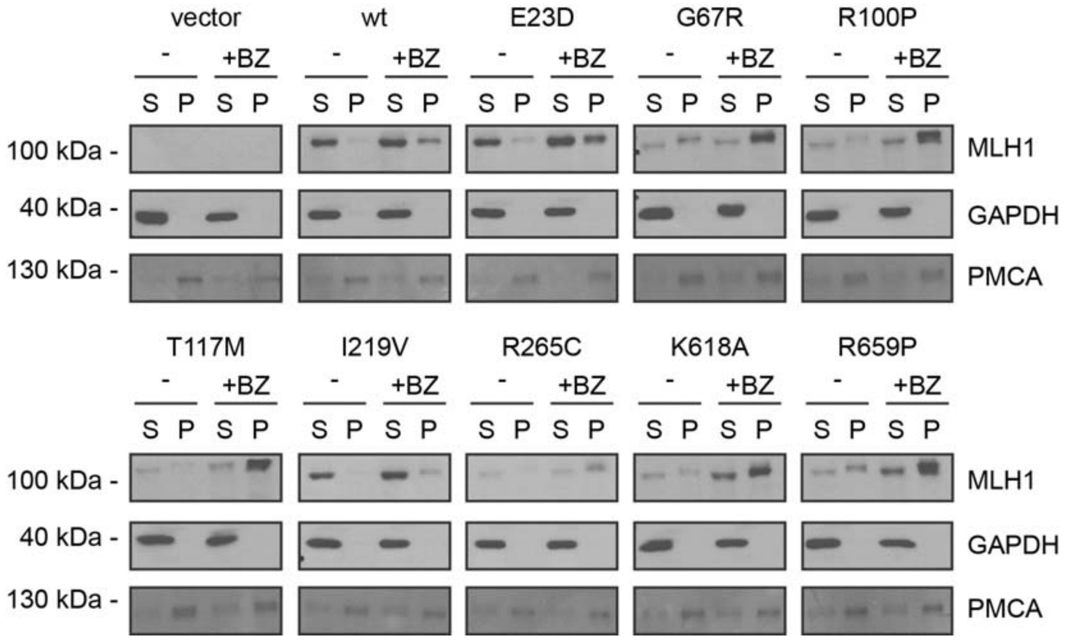


**Figure supplement – Figure 3** – *Solubility of selected MLH1 variants.* Transfected cells, either untreated (-) or treated for 16 hours with 10 μM bortezomib (+BZ) were lysed by sonication and separated into supernatant (S) and pellet (P) fractions by centrifugation. Western blotting using antibodies to MLH1 was employed to determine the amount of MLH1 in the fractions. Blotting with antibodies to GAPDH and PMCA served as loading controls for the soluble and insoluble fraction, respectively.

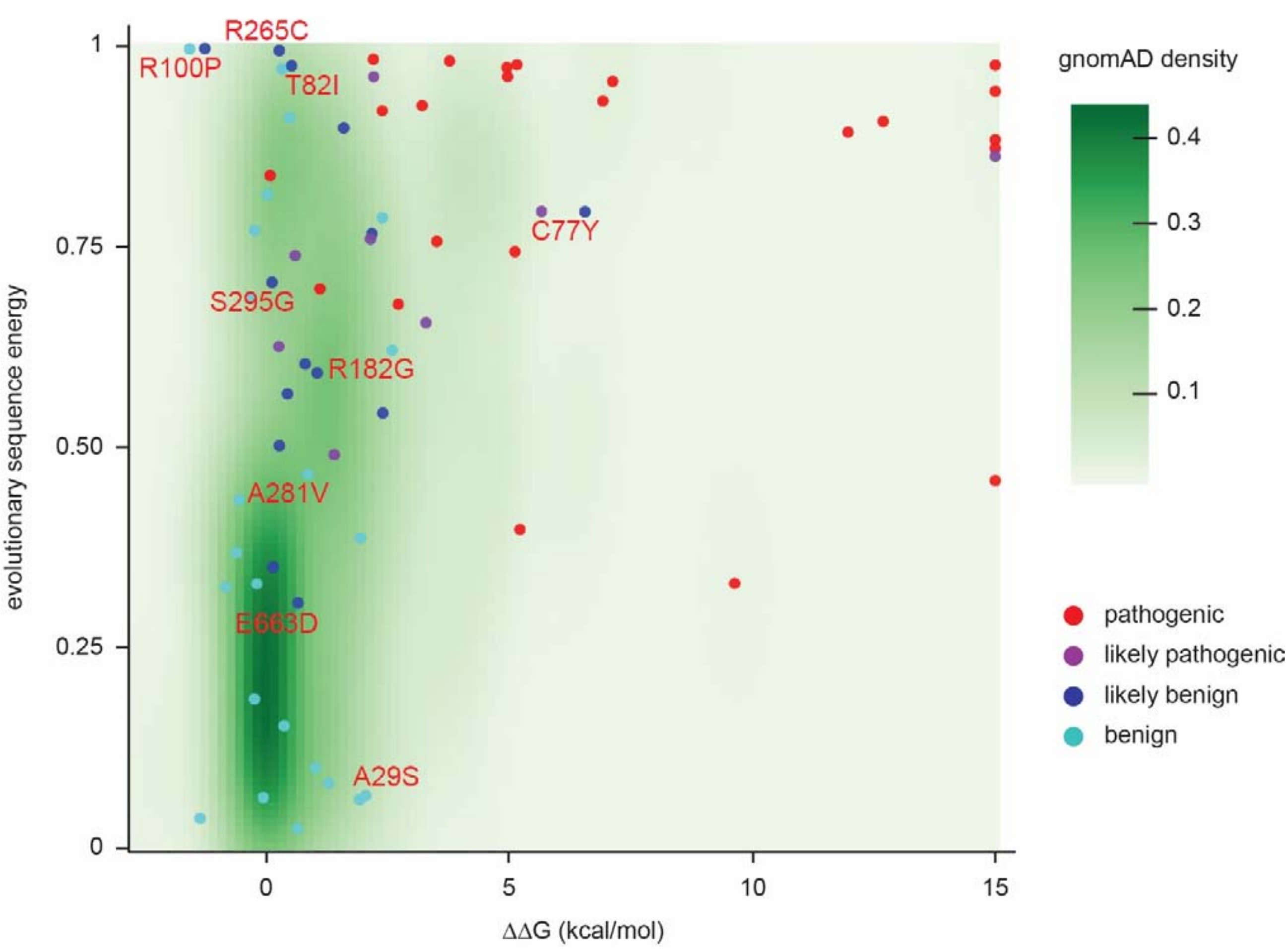


**Figure supplement - Figure 6.1** – *Landscape of ClinVar MLH1 variant tolerance*. Landscape of variant tolerance by combination of changes in protein stability (x axis) and evolutionary sequence energies (y axis), such that the upper right corner indicates most likely detrimental variants, while those in the lower left corner are predicted stable and observed in MLH1 homologs. The green background density illustrates the distribution of all variants listed in gnomAD. Outliers are discussed in the main text.

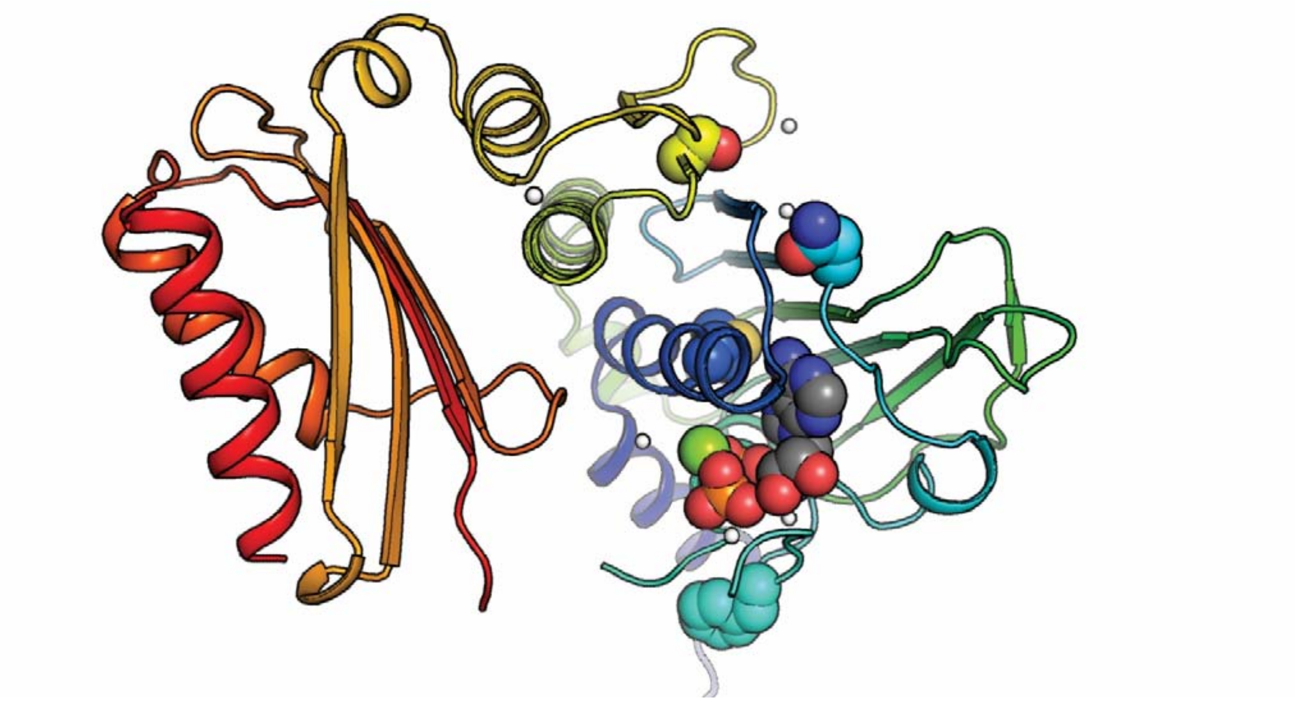

**Figure supplement – Figure 6.2 –** *Positioning of selected variants near the active site in the N-terminal domain*. The MLH1 structure (PDB: 4P7A) (Wu et al., 2015) with highlighted stable loss-of-function MLH1 variants (M35R, N64S, F80V, S193P) as spheres, as well as ADP. The domain is colored in a rainbow color scheme, with blue at the N-terminus and red at the C-terminal end of the N-terminal domain (MLH1 sequence position ~300); sidechain and ligand oxygen atoms are red, nitrogens blue, ligand carbons gray. Three of the stable loss-of-function positions (M35R, N64S, F80V) are very close to the ligand. Variation at these sites may thus interfere with ligand binding, which could explain why they lead to loss of function despite wild-type-like cellular protein levels.

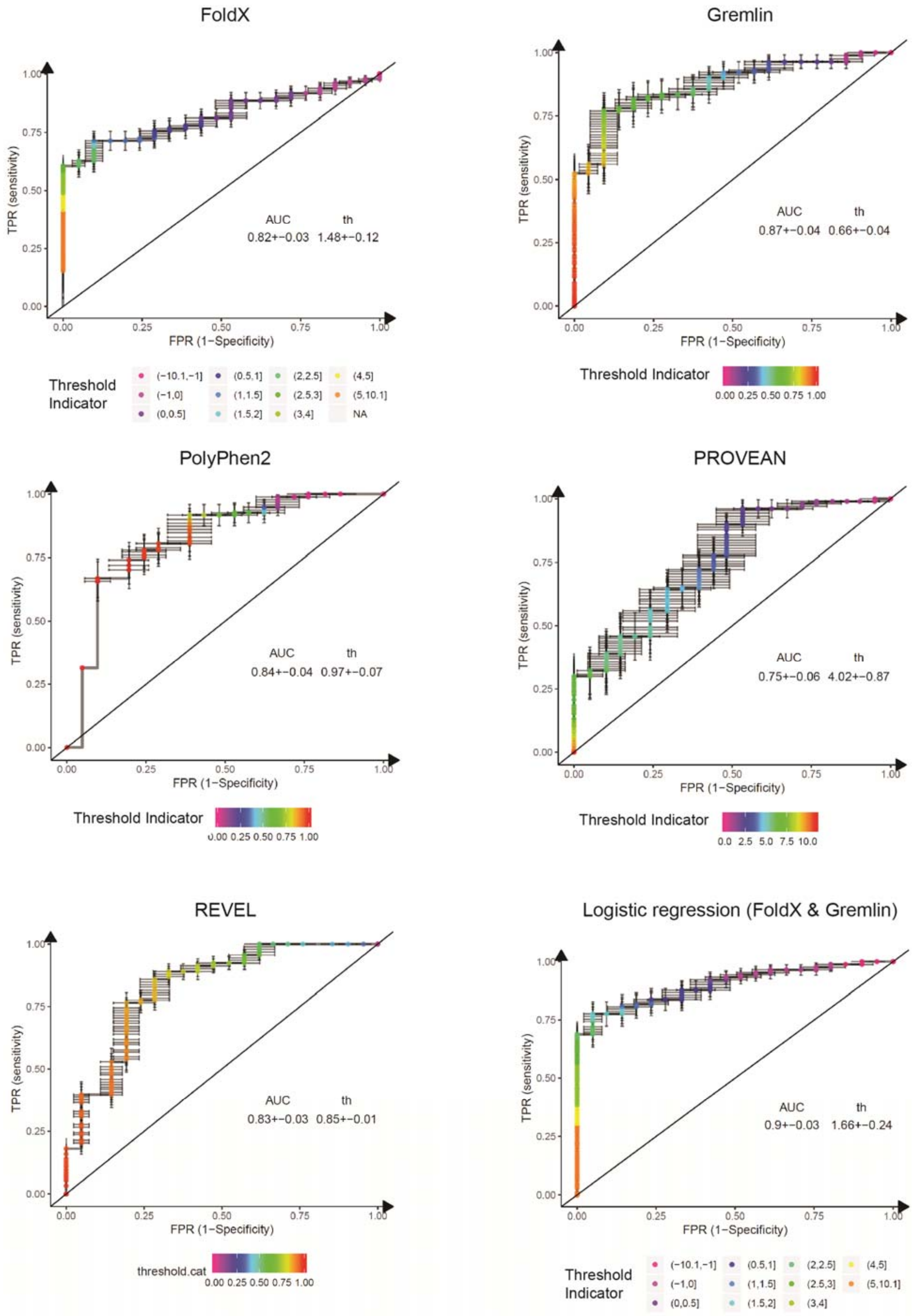
FoldX
TPR (sensitivity)
FPR (1−Specificity)
AUC th
0.82+−0.03 1.48+−0.12
Threshold Indicator
(−10.1,−1] (−1,0] (0,0.5] (0.5,1] (1,1.5] (1.5,2] (2,2.5] (2.5,3] (3,4] (4,5] (5,10.1] NA
Gremlin
TPR (sensitivity)
FPR (1−Specificity)
AUC th
0.87+−0.04 0.66+−0.04
Threshold Indicator
0.00 0.25 0.50 0.75 1.00
PolyPhen2
TPR (sensitivity)
FPR (1−Specificity)
AUC th
0.84+−0.04 0.97+−0.07
Threshold Indicator
0.00 0.25 0.50 0.75 1.00
PROVEAN
TPR (sensitivity)
FPR (1−Specificity)
AUC th
0.75+−0.06 4.02+−0.87
Threshold Indicator
0.0 2.5 5.0 7.5 10.0
REVEL
TPR (sensitivity)
FPR (1−Specificity)
AUC th
0.83+−0.03 0.85+−0.01
threshold.cat
0.00 0.25 0.50 0.75 1.00
Logistic regression (FoldX & Gremlin)
TPR (sensitivity)
FPR (1−Specificity)
AUC th
0.9+−0.03 1.66+−0.24
Threshold Indicator
(−10.1,−1] (−1,0] (0,0.5] (0.5,1] (1,1.5] (1.5,2] (2,2.5] (2.5,3] (3,4] (4,5] (5,10.1]

**Figure supplement – Figure 6.3** – *ROC curves for variant consequence predictors tested in this work.* The indicated predictors were used to score all benign, likely benign, likely pathogenic and pathogenic missense variants reported in ClinVar (Aug 2018). We merged benign and likely benign variants into the negative control set and pathogenic and likely pathogenic into the positive set and generated ROC curves for each predictor, assessing their power in separating benign from pathogenic variants. In addition to the individual predictors, we also assessed a logistic regression model combining FoldX $\Delta\Delta G$ and Gremlin evolutionary sequence energy ($\tilde{E}$). AUC, area under the curve. th, threshold (units specific to the respective predictor). Error bars indicate standard deviation from 100 randomly sub-sampled balanced ROC curves. See also Fig. 6E in the main manuscript.
